# Classification and Regression Trees to predict Transcription Factor Combinatorial Interaction in scRNA-seq data

**DOI:** 10.1101/2024.04.17.589552

**Authors:** Jean Baptiste Carluer, Laura Steinmann, Clément Carré, André Mas, Gabriel Krouk

## Abstract

Understanding the regulatory mechanisms that govern gene expression is crucial for deciphering cellular functions. Transcription factors (TFs) play a key role in regulating gene expression. In particular TF combinatorial interactions (TFCI) are now thought to largely shape genomic transcriptional responses, but predicting TFCI *per se* is still a difficult task. Single-cell RNA sequencing (scRNA-seq) has emerged as a powerful tool providing a whole new readout of gene regulatory effects. In this study, we propose a machine learning approach utilizing Classification and Regression Trees (CART) for predicting TFCI in >110k scRNA-seq data points yielded from *Arabidopsis thaliana* root. The proposed methodology provides a valuable tool for pointing to new TFCI mechanisms and could advance our understanding of Gene Regulatory Networks’ functioning.

## Introduction

Transcription factors (TFs) are orchestrating the cell regulatory landscape by binding to their cognate DNA motif (Blanc-Mathieu et al., 2022). The famous ABC model that defines plant flower development demonstrated that TF combinatorial games are crucial to trigger different developmental genetic programs (Bowman et al., 1991). This model states for instance that the combination of 2 types of TF is necessary to trigger the development of appropriate flower organs such as stamen or petals. More recently, TF combinatorial interactions (TFCI) have been explained at a structural level. For instance LEAFY (LFY), being a major regulator of flower development, changes its binding site preference under the cooperative action of UNUSUAL FLORAL ORGANS (UFO) (Rieu et al., 2023). Another recent example shows that bZIP transcription factors are in combination binding to variable Cis-Regulatory-Elements (CREs) (Li et al., 2023). From a regulatory perspective it has also been demonstrated that genome wide transcriptomic response triggered by Nitrogen provision is also the result of a combinatorial action of TFs (Brooks et al., 2019). Thus literature now shows that TFCI are a very important phenomenon that might be a general rule rather than an exception.

scRNA-seq are on the path to change transcriptomic studies forever as they provide an unprecedented accurate measure of gene expression in every cell of an organism and a wealth of data (Shahan et al., 2022; Nolan et al., 2023). Among the first scRNA-seq studies, we count those applied to Arabidopsis root being a very good model for such studies thanks to its concentric well defined cell layers that recapitulate a very well defined developmental program (Shahan et al., 2022). scRNA-seq techniques yield several thousand data points per experiment. As a matter of dimension in data acquisition, the first few papers reporting Arabidopsis scRNA-seq studies (Denyer et al., 2019; Jean-Baptiste et al., 2019; Ryu et al., 2019; Shulse et al., 2019; Zhang et al., 2019) yielded > 110k data points (cells) that directly overpassed the totality of RNA-seq data points in GEO repository at that time. Now nuclei isolation and RNA sequencing can reach millions of data points (Lee et al., 2023; Nobori et al., 2023). Despite this huge advance in data acquisition, the data structure of scRNA comes with drawbacks. Indeed, the depth of sequence and the number of genes being sequenced per cell/nuclei is rather limited (in the order of a couple thousand genes). This noisy characteristic of scRNA-seq data can be explained by mainly two reasons. First, technically the amount of RNA being retrieved per cell is too limited to get a complete survey of the actual transcripts present in the cell. Second, it might be a biological characteristic that gene expression is inherently noisy and that each cell does not necessarily express all the repertoire of genes. These two points are not mutually exclusive and are a matter of debate (Sanchez and Golding, 2013; Jovic et al., 2022).

Gene Regulatory Networks are constructed in many different ways and are data greedy (Krouk et al., 2013; Van den Broeck et al., 2020). One particularly interesting approach used to detect TF interaction is the use of linear models either in the form of simple linear systems or in the form of ODEs. These attempt to fit a given gene expression as a linear composition of regulatory variables (most often TF expression) (Ristova et al., 2016; Krouk et al., 2013; Van den Broeck et al., 2020). However these kinds of models often struggle with sparse and noisy data such as scRNA-seq.

In this work we established a data analysis pipeline for detecting TFCI events in large scRNA-seq data. We used a machine learning technique named CART to perform this task embedded in a whole suite allowing direct visual evaluation of the model outcome.

## Results

In an attempt to still exploit the wealth of scRNA-seq and detect TFCI, we built a dedicated bioinformatic pipeline named CART-CELLS whose scheme is described in Figure 1. CART-CELLS is a comprehensive approach that integrates expression profiles obtained from single-cell data with TF expression to infer TFCI centered gene regulatory networks in a user-friendly manner.

**Figure 1.**
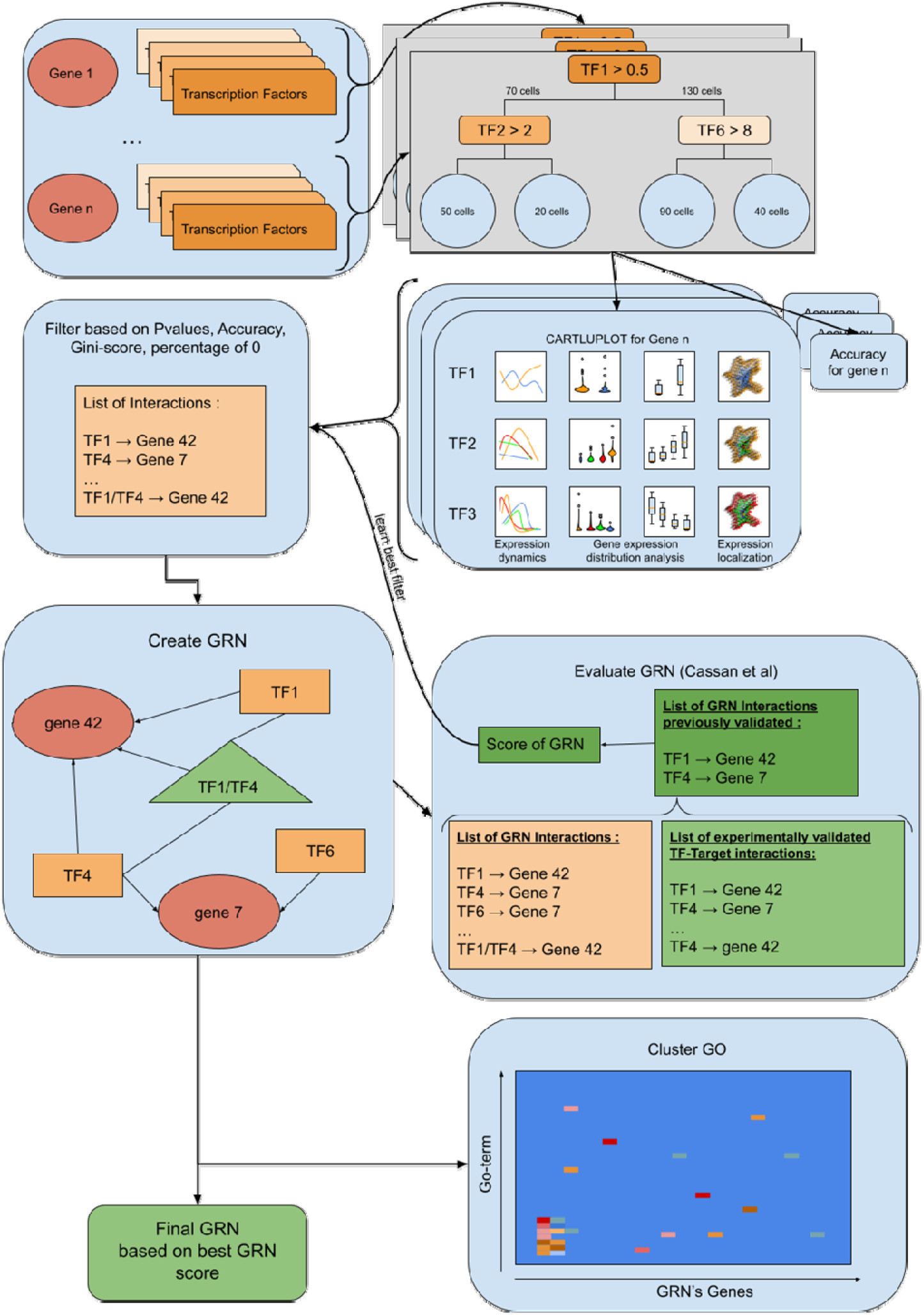
scRNA-seq processing rationale to reveal TFCI. The first step is the generation of the CART tree and its evaluation. Cart-tree results analysis is plotted and compiled. The second step is the generation of GRN according to CART tree results (provide Sup Table 1). Filter will be learned using Cassan et al (2023) evaluation. Once the best model is found the final GRN is processed and according to actors the cluster GO plot will be generated.

### Using CART to find TFCI

The core of our algorithm is Classification and Regression Trees or CART (Gordon et al., 1984). It is a non-parametric machine learning technique that recursively partitions the feature space (here cells) based on the most informative variables (here TF). In other words, a given gene expression across the 110k cells is partitioned in classes for which a given TF reaches a given threshold (α) or not.

The proposed method can be divided into three main components, with an optional fourth, for inferring TFCI enhanced GRNs from single-cell RNA seq data (Figure 1).

Firstly, decision trees are constructed for each single gene in the genome having a non null expression in the dataset (here 24 958 genes), from which decision rules will be extracted (Figure 1). For each target gene (TG) three TFs (TF1, TF2, TF3) are selected by CART as being the best explanatory variables with a certain threshold (α1, α2, α3 respectively) of TF expression.

### CARTLU-Plot to visualize the best TFCI

From this, for each TG a 3 layer plot is generated named CARTLUPLOT Figure 1 (scheme) and Figure 2 (real data). The utilization of these representations enables us to visually analyze a substantial amount of information, facilitating the evaluation of whether the transcription factors are a potential candidate for the regulation of the TG alone (first level) or in combinations (second and third level).

**Figure 2.**
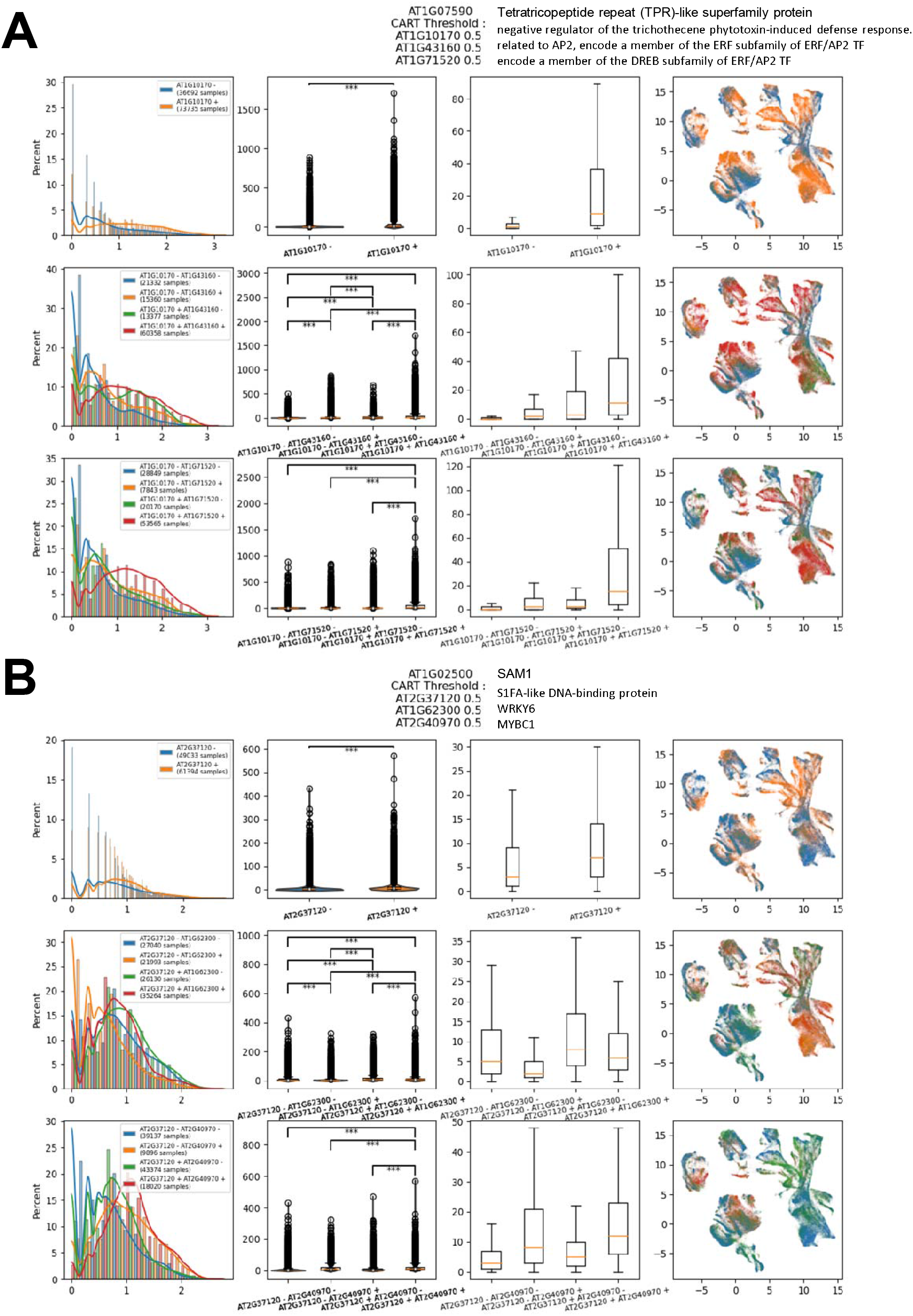
CART LEVEL-UP PLOT (CARTLU-PLOT) CART procedure leads to identification of potential TFCI for two target genes : AT1G07590 (A) and AT1G02500 (B). Représentation scheme is the same for each of them. The first row focuses on TG-TF1 interaction while the second and third row focus on the interaction between TG and (TF1-TF2) and between TG and (TF1-TF3) respectively. First column is a representation of the expression’s dynamics using curves and line plots. To assess good representation while the dimorphism between conditions (e.g. TF1+ and TF1-) is important, a normalization has been made. The second and third columns are representations of the gene expression distribution using boxplot, the only difference between these two plots is the extraction in the third plot of the outlier using the “showfliers” parameters from matplotlib. The last column is a UMAP representation, which illustrates any dimorphism resulting from the condition (particular TF combination) and may help to determine if specific cell types are targeted by the mechanism.

In detail, for each TG (having a non null transcriptomic signal) we designate the TF that is predicted by CART to have the most influence on a given TG as TF1.The data for this given gene is then parsed between cells for which TF1 is > α1 that and cells for which TF1 is < α1. The initial plot (top row Figure 2) depicts the expression distribution of the TG under two TF1 conditions: TF1>α1 (referred as TF1+) and TF1<=α2 (referred as TF1-). This allows us to visualize the potential influence of TF1 on TG (Figure 1 and Figure 2).

Next, the data is parsed to visualize the combinatorial influence of TF1 and TF2 (Figure 2, line of plots #2). The data for these given genes is then parsed in 4 batches. The first batch contains cells for which TF1 is < α1 and TF2 < α2 (referred as TF1-, TF2-). The second batch contains cells for which TF1 is < α1 and TF2 > α2 (referred as TF1-, TF2+). The third batch contains cells for which TF1 is > α1 and TF2 < α2 (referred as TF1+, TF2-). The fourth batch contains cells for which TF1 is > α1 and TF2 > α2 (referred as TF1+, TF2+). The following plot depicts the expression distribution of the TG under these 4 conditions of TF combination. This allows us to visualize the potential influence of TF1 and TF2 combinatorial influence on TG (Figure 2). Finally the same operation is applied to TF1 and TF3 combinatorial influence.

The following plots (second row Figure 2) depicts the expression distribution of the TG under four conditions (TF1-,TF2-), (TF1+,TF2-), (TF1-,TF2+), (TF1+,TF2+). On the same line the following plots use outlier removal technique (using the “showfliers” parameter from matplotlib’s boxplot). These plots primarily aim to examine the mean, quantile, and extremum values for each case. We then use non-parametric Mann-Whitney statistical tests to compare the distributions under each condition. The last representation on the line is a Uniform Manifold Approximation and Projection (UMAP) analysis performed on the complete dataset. This analysis illustrates any dimorphism resulting from the condition (particular TF combination) and may help to determine if specific cell types are targeted by the mechanism. UMAP has already been used for gene expression analysis according to cell types (Nolan et al., 2023).

### Building Gene regulatory network with TFCI influence

Gene regulatory network plays a fundamental role in orchestrating the complex regulatory processes that govern gene expression and cellular behavior. These networks consist of interconnected nodes representing genes and regulatory elements.

Understanding the structure and the dynamics of GRNs is crucial for unraveling the intricate mechanisms underlying cellular processes, development, and diseases. Here our method builds upon TFCI interaction detection by CART in order to implement the best ones in a GRN structure.

A GRN depicts the intricate web of interaction between genes and regulatory elements (here TF), governing the activation or repression of gene expression (Figure 3).

**Figure 3.**
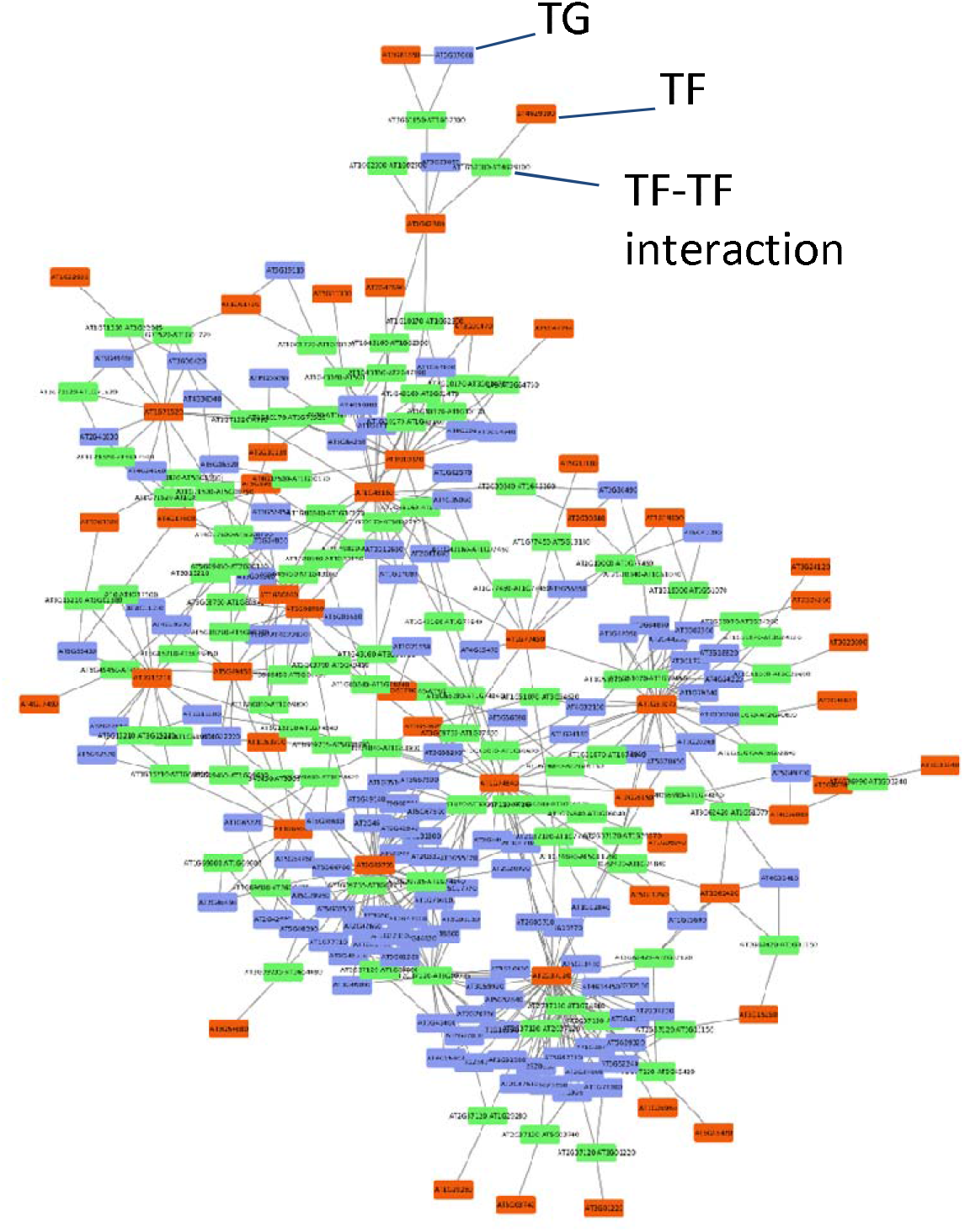
Cytoscape view of an Inferred gene regulatory network (GRN) containing TFCI. The parameter selection procedure has allowed the selection of this model with the following parameters : -log10(pval) superior or equal to 50, accuracy superior or equal to 0.3, percentage of zero superior to 30%, gini score superior or equal to 0.9. Nodes represent the genes, TF or TF interaction based on colors, green represents TF interactions, blue represents TGs and red represents TFs. There is 51 TF, some of them are isolated and are found in only one TF-TF interaction (e.g. AT1G22985, AT4G29100,…), some other are central node and then highly connected to other nodes (e.g. AT2G37120, AT3G09735, …) and some of them are apparently highly connected to TG and found in direct connection (TF-TG interaction) as well as in indirect connection (TF1-TF2 interaction with TG). Next, there are 103 TF-TF interactions and 123 TG involved in this GRN.

A GRN can be conceptualized as a complex system of interconnected components, akin to a dynamic network where information flows through regulatory interactions. At the center of this network lie TFs. TFs can activate or repress target genes, either directly or indirectly, through a series of intricate interactions with other components of the network. GRN structure is typically represented as a graph denoted as *G =* (*V, E*), with nodes also called vertices *V* and links also called edges *E* that capture regulatory interactions between genes and regulatory elements.

In a GRN, a node represents a gene or regulatory element involved in gene regulation. Here we implement a new kind of node representing the potential combinatorial influence of two TFs based on the CART modeling described above (Figure 3).

***Nodes*** now represent three potential elements: TG, regulatory elements (here TF), and interactions between two transcription factors (aka TFCI). By adopting this representation, we were able to focus our investigation on the interactions between transcription factors and identify which combinations of transcription factors are more frequently observed and thus potentially important at the genome level.

***Edges*** are inherently directed from a TF or TFCI to a gene. It is also evident that TF nodes can be connected to an interaction of TF nodes, with the directionality of the connection from the TF to the interaction node. This network representation exhibits biological parallels, where two individual transcription factors can for instance physically interact or compete for a binding site, forming connections or exerting influence on each other.

### Parameter filtering to build GRN with TFCI influence

In order to select the optimal sub-population of gene regulatory networks (GRNs), a carefully designed pipeline has been established to filter GRN nodes based on various parameters being CART model accuracy, CART related Gini score and, percentage of 0 in TG expression vector, *p value* returned by Mann Witney tests. Undoubtedly, one might assume that high model accuracy or a favorable Gini score serve as reliable indicators. However, it’s important to acknowledge that these metrics are influenced by the percentage of 0 in TG expression, introducing a bias. As a result, determining the optimal parameters is not as straightforward as initially perceived.

Multiple values have been thoroughly examined for each of the four parameters and their exhaustive combination. This represents 420 different GRNs that have been assessed with each of the 4950 combination of parameters below (Sup Table 1):

- CART model accuracy: 0.0, 0.1, 0.2, 0.3, 0.4, 0.5, 0.6, 0.7, 0.8, 0.9, 1.0

- Gini score: 0.1, 0.2, 0.3, 0.4, 0.5, 0.6, 0.7, 0.8, 0.9

- Percentage of 0 in TG expression: 0.5, 0.4, 0.3, 0.2, 0.1

-log10(Mann-Whitney test *p value*): 50, 40, 30, 20, 15, 10, 8, 5, 3, 2

The underlying concept is to apply each filter to the data (matrix containing for each TG returned parameters [Sup Table 1]) and subsequently assess the resulting GRN for instance by following (Cassan et al., 2023) procedure. Subsequently, by analyzing the ratio between the number of target genes in the GRN and the evaluation of the GRN, optimal parameters can be determined (Figure S1).

The executed protocol resulted in the generation of two Gene Regulatory Networks (GRNs), as illustrated in Figure 3 and Figure S2, which exhibit the optimal recall-to-node-number ratio. Intriguingly, these two GRNs manifest conserved functionalities, as revealed by the Gene Ontology (GO) analysis presented subsequently, where specific functions are found to be influenced by common transcription factors (TFs). This observation is indicative of a non-random genesis of GRNs (Krouk et al., 2013; Medici et al., 2015).

### Cluster GO map

Indeed, most often the quality of GRN cluster and TF-TG influence can be measured by the coherence Cluster GO map. It serves as a concise depiction of the GO term associated with each transcription factor discovered within the final GRN. In conducting this analysis on GRN presented Figure S4, we search for transcription factors within the GRN and retain only the first-order target nodes. Subsequently, we record and examine each GO term linked to the respective TG present in the target nodes within the local group. The purpose of this analysis is to determine whether a specific transcription factor has been consistently associated with a particular function more frequently than expected by chance. Such findings would likely carry greater significance in the final analysis of the TF-TG connection.

### Biological insights

Hereby we would like to highlight a few biological insights that can be revealed by our analysis pipeline. Figure 2A presents the stunning combinatorial regulation of AT1G07590. This gene codes a Tetraticopeptide repeat (TPR) -like superfamily protein. The three TF that have been identified as having an effect on the TG are: AT1G10170, AT1G43160 and AT1G71520. They respectively code for a negative regulator of trichothecene phytotoxin-induced defense response, a gene related to AP2 that encode a member of the ERF subfamily of ERF/AP2 TF and a gene that encode a member of the DREB subfamily of ERF/AP2 TF.

In the presence of AT1G10170 the target gene appears to be dramatically more expressed (Figure 2A, top row). More importantly, the co-occurence of AT1G43160 and AT1G10170 synergistically enhance the production of the TG (Figure 2A, second row). Conversely, when both AT1G43160 and AT1G10170 are absent, TG production remains markedly reduced. A similar pattern of synergistic behavior is observed in the interaction between AT1G71520 and AT1G10170.

Another example of potential antagonistic effect can be reported for AT1G02500 as TG (Figure 2B). AT1G02500 is a gene that encodes SAM1 being a S-adenosylmethionine synthetase. In the presence of AT2G37120, the TG expression tends to be moderately enhanced. The interesting point is the interaction of AT2G37120 with AT1G62300. Indeed, we can observe that the potential positive effect of AT2G37120 on the TG is likely counteracted when AT1G62300 is present. Thus defining a potential antagonistic effect. On the other hand, AT2G37120 displays a synergistic action on the TG as the presence of both TF enhances the TG expression more than when both are expressed alone.

## Discussion

In conclusion, with our procedure, each of the >110,000 cells happens to be a particular witness of the potential combinatorial effect of TFs. We believe that this work may provide a valuable suite of tools to infer TFCI in single cell RNAseq data that are inherently noisy and sparse. This has been made possible by the use of the CART algorithm combined with powerful visualization tools custom built to display each gene in the genome. However this method is limited by two main points: Firstly, factual validation by external dataset is rather limited, reaching pretty low recall (Figure S1). This can easily be explained by the fact that TF-TG databases are still limited as compared to the world of potential interactions between TFs and TGs. It is worth noting here that if the connections that we are revealing here have not been shown experimentally before, maybe because of their inherent nature of tissue specific expression, low recall value does mean bad results. For instance by analyzing the GRN Figure S2, which is the GRN with more flexible parameters, we were able to see that for 946 interactions in the GRN only 15 of them have been found by CHIPSeq, 61 of them by DAPSeq, 619 of them are not studied and 251 seems not to be supported. Secondly our method is now adapted and mainly dedicated to detect TFCI. To our knowledge there is currently no publicly available database large enough susceptible to validate these new discoveries. This would be a valuable experimental resource that is currently under development in our lab.

## Conclusion

This method/algorithmic package provides a valuable tool to understand genetic regulation and shows that CART based approach on scRNA-seq is full of promise to predict TFCI. We hope that future work in this domain could lead to high throughput validation and then allow our method to show his full potential. These research could bring important commitment to solve genetic regulation mechanisms in the cell and then could pave the way to new discoveries on genetic regulatory networks in plants and potentially other model organisms.

### Methods Dataset

We used the dataset previously described in (Shahan et al., 2022). The whole normalization procedure has been repeated and yielded in our hands similar results. It generated the matrix M containing i columns and n rows corresponding respectively to 110,427 cells representing all major cell types and 24,997 genes This matrix is available on NCBI (https://www.ncbi.nlm.nih.gov/geo/query/acc.cgi?acc=GSE152766).

### Code

The whole code is provided on https://github.com/CarluerJB/CART_CELLS for anyone to use under open source license.

## Author contributions

Conceptualization: G.K, JB.C, A.M Methodology: A.M, J.B.C. (Statistical Analysis, Machine Learning) Writing – original draft: G.K, JB.C, A.M

## Supplementary Materials

**Sup Table 1: Classification and Regression Tree results for each gene**.

Model score describe the global strength of the model while gini score describe the impurity of each found condition, TFs conditional acting on the TF are specified if found (e.g. TF1<=17.5) Mann Whitney p-value are given for each condition as well as population structure information (mean for each condition, percentage of zeros).

**Sup Table 2: GRN evaluation results for each parameter**.

P-value, percentage of zero, model score and gini score variation allow to create new GRN, GRN information are indicated (nb_gene, nb_nodes, nb_edges), then GRNs are evaluated using Cassan evaluation method (AranetBench), which allow to have quantify interaction that have already been found in literature.

**Sup Info:** GRNs from Figure 3, and Figure S2 are provided as .cys files.

---

**Figure S1.**
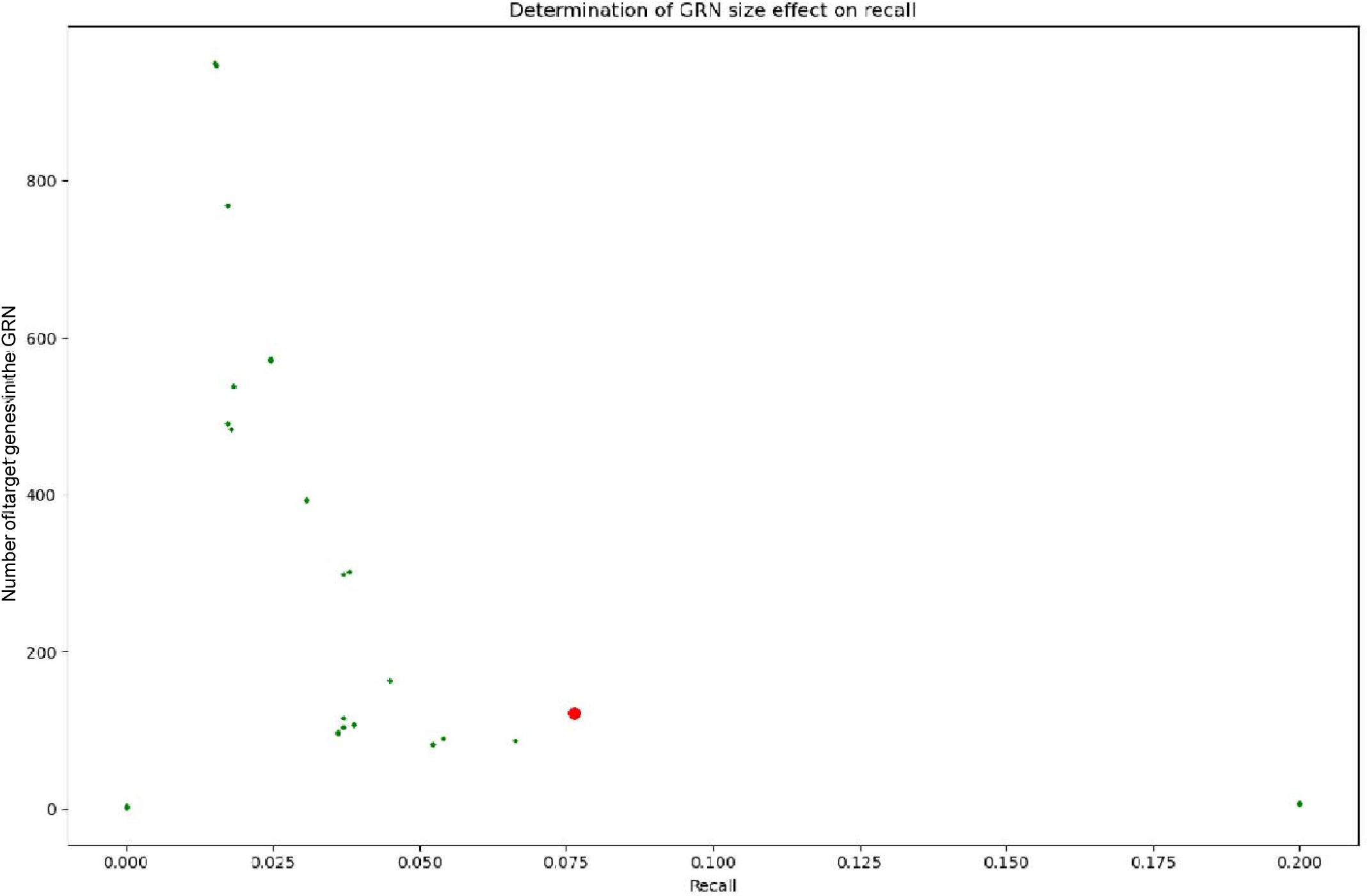
Performance analysis of parameter selection based GRN models. The parameter selection procedure ends with model evaluation using Cassan et al 2023 GRN evaluation method, the returned recall is used in this scatter plot and compared with the number of target genes in each GRN. Recall goes from the lowest 0 to 0.2, while GRN size goes from 0 to less than a thousand of nodes. These low recall values are discussed in the text.

**Figure S2.**
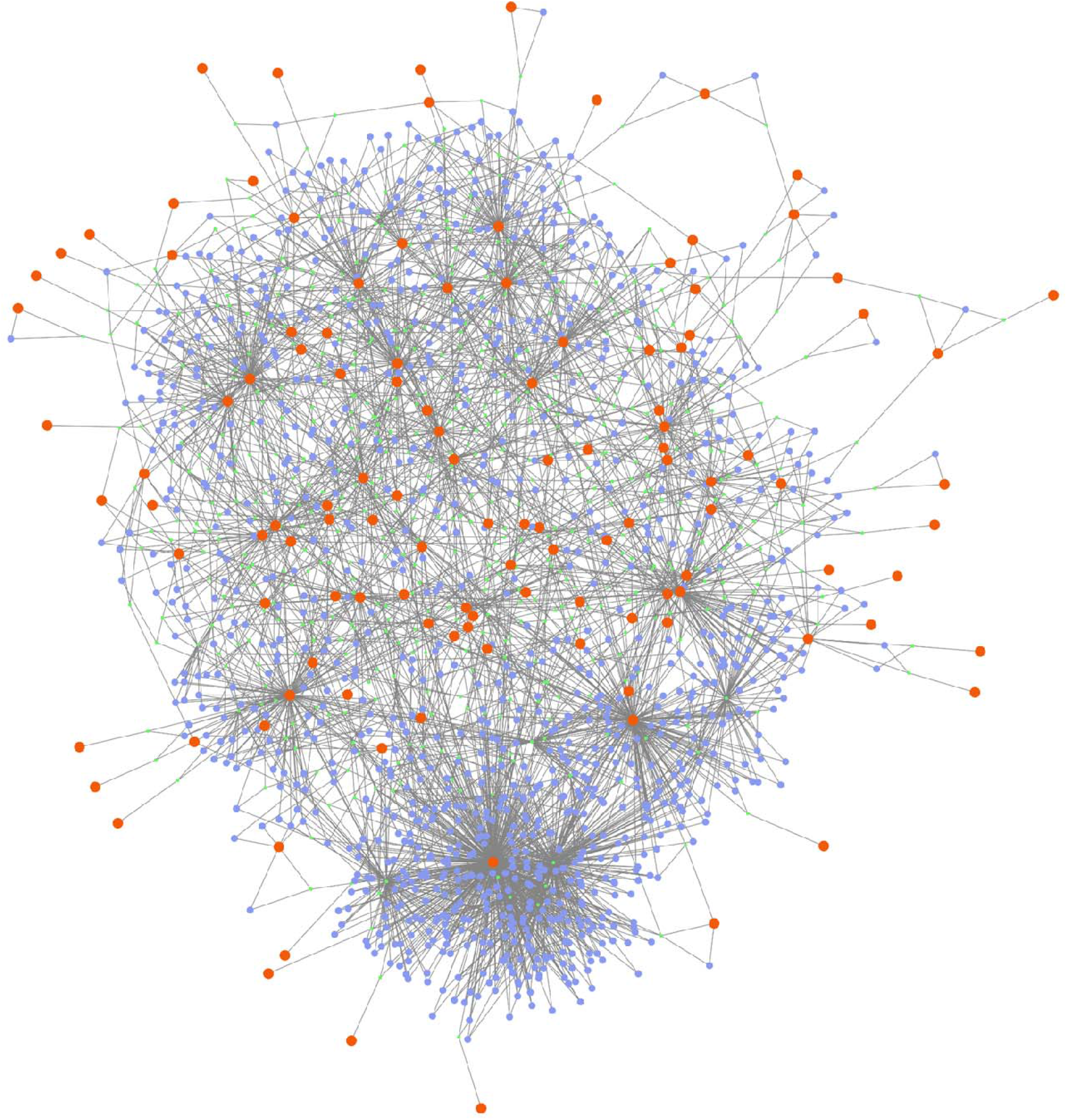
Cytoscape view of an Inferred gene regulatory network (GRN) based on relaxed parameters. These parameters only focus on the percentage of total zero (superior to 50%) and on the -log10(pval) superior or equal to 20 for the Mann Whitney tests. The network is more dense than the network from parameter selection; there are 949 TG nodes, 518 TF-TF interaction and 117 TF involved in the GRN.

**Figure S3.**
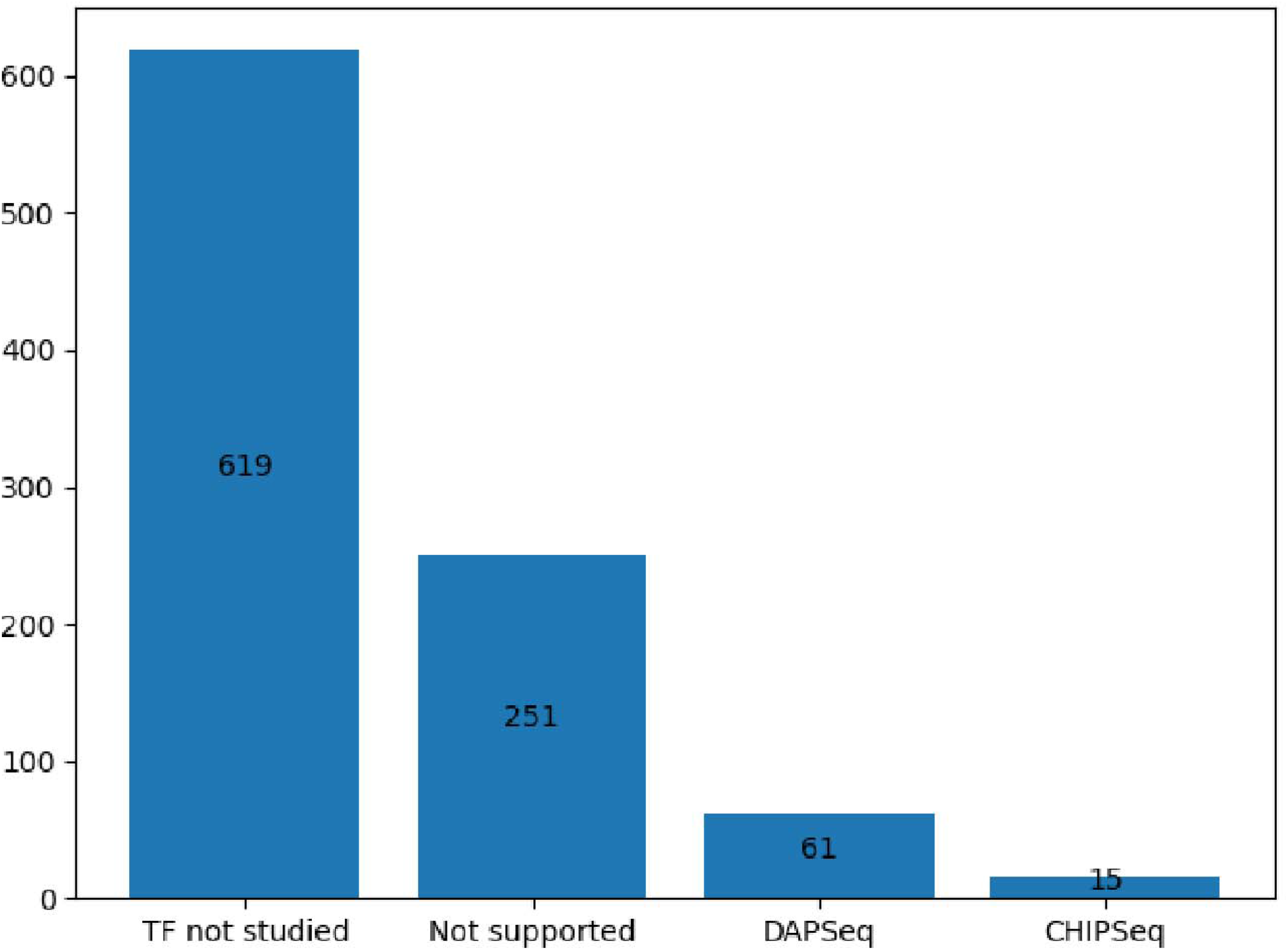
GRN validation method results. Among the 946 interactions, 619 are linked to unstudied transcription factors, 251 not supported transcription factors, 61 linked to DAPSeq validated interactions and 15 to CHIPSeq validated interactions

**Fig. S4.**
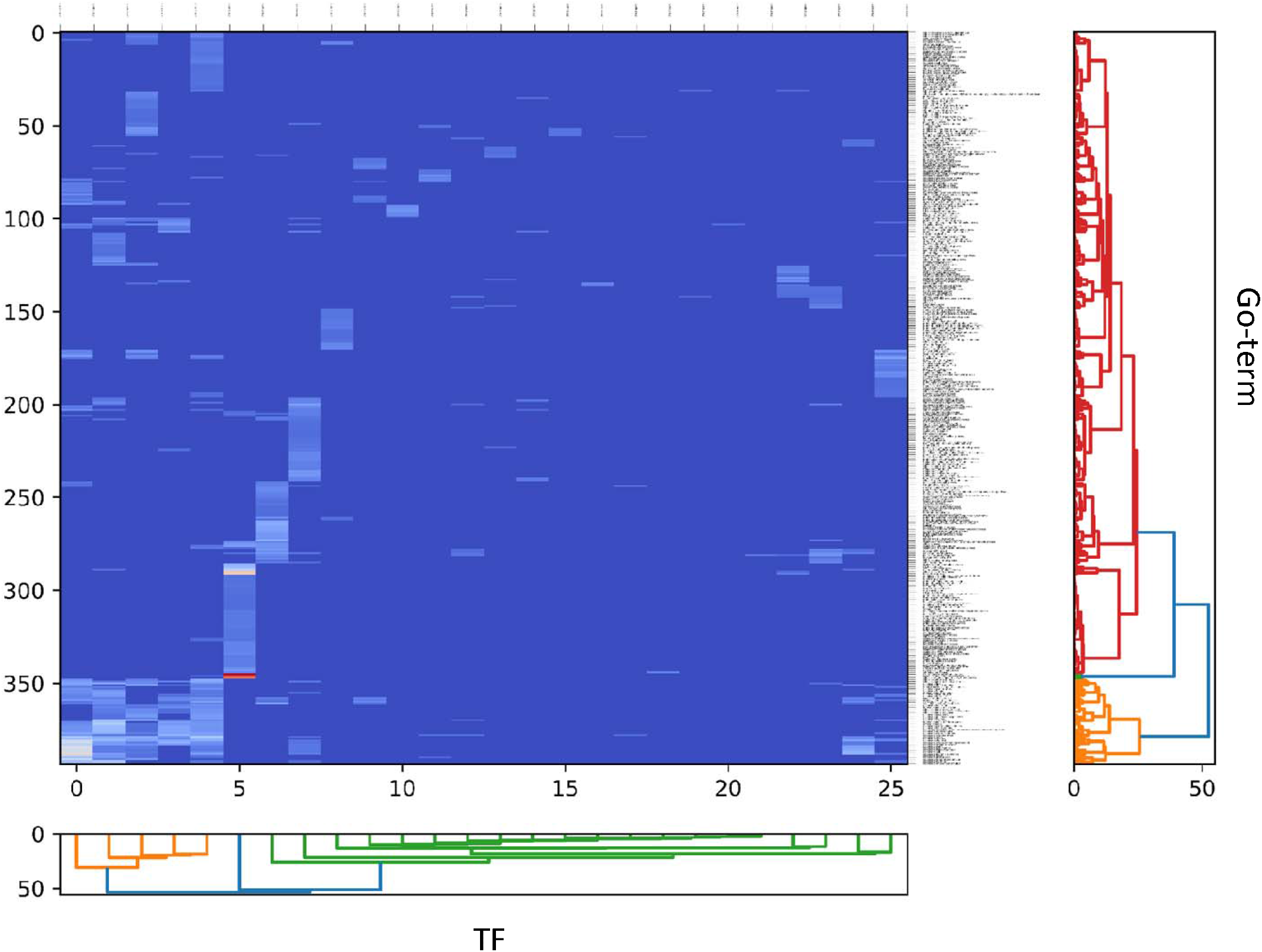
Cluster Go map, dendrogram of Go-term and Transcription Factors. Heatmap that represent Go term on the y axis and Transcription Factors on the x axis, each axis has been clusterized according to correlation score between the association of TF with specific GO-term. To compute this score the GOATOOLS platform has been used to identify non random connections between TF and TG. The metric used to evaluate this correlation is the p_uncorrected with a p-val fixed at 0.05. The most important signal come from the TF AT1G74840, which is identified to nitrogen and macromolecule linked process (cellular nitrogen compound biosynthetic process, organonitrogen compound biosynthetic process, macromolecule biosynthetic process).

